# Glucose-1,6-bisphosphate, a key metabolic regulator, is synthesized by a distinct family of α-phosphohexomutases widely distributed in prokaryotes

**DOI:** 10.1101/2021.12.02.470922

**Authors:** Niels Neumann, Simon Friz, Karl Forchhammer

**Affiliations:** Interfaculty Institute of Microbiology and Infection Medicine, University of Tübingen, Auf der Morgenstelle 28, 72076 Tübingen, Germany; Cluster of Excellence: EXC 2124: Controlling Microbes to Fight Infection, Tübingen, Germany

**Keywords:** Phosphoglucomutase, glucose-1,6-bisphosphate, fructose-1,6-bisphosphate, glycolysis, carbon metabolism

## Abstract

The reactions of α-D-phosphohexomutases (αPHM) are ubiquitous, key to primary metabolism and essential for several processes in all domains of life. The functionality of these enzymes relies on an initial auto-phosphorylation step which requires the presence of α-D-glucose-1,6-bisphosphate (Glc-1,6-BP). While well investigated in vertebrates, the origin of this activator compound in bacteria is unknown. Here we show that the Slr1334 protein from the unicellular cyanobacterium *Synechocysitis* sp. PCC 6803 is a Glc-1,6-BP-synthase. Biochemical analysis revealed that Slr1334 efficiently converts fructose-1,6-bisphosphate (Frc-1,6-BP) and α-D-glucose-1-phosphate/α-D-glucose-6-phosphate into Glc-1,6-BP and also catalyzes the reverse reaction. As inferred from phylogenetic analysis, the *slr1334* product belongs to a primordial subfamily of αPHMs that is present especially in deeply branching bacteria and also includes human commensals and pathogens. Remarkably, the homologue of Slr1334 in the human gut bacterium *Bacteroides salyersiae* catalyzes the same reaction, suggesting a conserved and essential role for the members of this αPHM subfamily.

## Introduction

The α-D-phosphohexomutase (αPHM) superfamily is ubiquitous and found in all domains of life. All known members of this superfamily catalyze a reversible intramolecular phosphoryl transfer on their phosphosugar substrates. This reaction requires a bound metal ion (usually Mg^2+^) together with a conserved phosphorylated seryl residue in the active site and proceeds via a bis-phosphorylated sugar intermediate ^1^. While showing similar properties in structure and reaction mechanism, these enzymes differ strongly in substrate specificity. Based on their preferred substrate, the αPHMs are traditionally divided into four main groups: (1) Phosphoglucomutases (PGM), (2) phosphoglucosamine mutases (PNGM), (3) phosphoacetylglucosamine mutases (PAGM) and (4), the group of bifunctional phosphoglucomutases/phosphomannomutases (PGM/PMM) ^1^. The most recent expansion to this classification has been the discovery of the mammalian phosphopentomutase (PGM2) and Glucose-1,6-Bisphosphate-synthase (PGM2L1) ^2, 3^. This classification, however, lacks a more detailed subdivision in between the main groups and also does not take into account αPHMs preferably catalyzing other reactions (e.g. straight phosphomannomutases) ^3^. The NCBI conserved domain database (CDD), instead, classifies the αPHM superfamily in 11 different subfamilies on a phylogenetic basis ^4^. This phylogenetic model enables a more detailed classification and includes several groups of bacterial and archaeal enzymes showing typical αPHM properties but being only poorly characterized (**Table 1**).

**Table 1:** Subfamilies in the αPHM superfamily with their identifier and proposed main function according to the NCBI conserved domain database (CDD)

| CDD group name | CDD identifier | Proposed main function |
| --- | --- | --- |
| PGM1 | cd03085 | Phosphoglucomutase |
| PGM2 | cd05799 | Phosphopentomutase (PGM2)/Glc-1,6-BP synthase (PGM2L1) |
| PGM3(PAGM) | cd03086 | Phosphoacetylglucosamine mutase |
| PMM/PGM | cd03089 | Bi-functional phosphomanno-/phosphoglucomutase |
| PGM like 1 | cd03087 | Unknown |
| PGM like 2 | cd05800 | Unknown |
| PGM like 3 | cd05801 | Phosphoglucomutase |
| PGM like 4 | cd05803 | Unknown |
| ManB | cd03088 | Phosphomannomutase |
| GlmM | cd05802 | Phosphoglucosamine mutase |
| Mpg1 transferase | cd05805 | GDP-mannose-pyrophosphorylase |

The PGM (cd03085) from rabbit and human as well as the bacterial PGM (cd05801) from *Salmonella typhimurium* and the bacterial bifunctional PMM/PGM (cd03089) from *Pseudomonas aeruginosa* are among the best characterized αPHMs with available crystal structures and detailed description of reaction mechanisms ^5-9^. Catalyzing the interconversion of α-D-glucose-1P (Glc-1P) and α-D-glucose-6P (Glc-6P) makes PGM a key enzyme in glycogen metabolism, glycolysis and gluconeogenesis. During this reaction, a phosphoserine at the active site of PGM donates the phosphoryl group to the substrate Glc-6P or Glc-1P resulting in the synthesis of the transient intermediate α-D-glucose-1,6-bisphosphate (Glc-1,6-BP). After a reorientation in the catalytic center, the intermediate product re-phosphorylates the active site serine yielding Glc-1P or Glc-6P, respectively. Glc-1,6-BP itself acts as an essential activator of PGM by providing an initial phosphorylation of the active site and was first discovered by Leloir, Trucco ^10^ in yeast extract ^11^. While Glc-1,6-BP is not a free metabolite in any metabolic pathway, it is known to be a potent regulator of several enzymes in central carbon metabolism in eukaryotes: in addition to PGMs and other αPHMs, it was shown to regulate hexokinases, 6-phosphogluconate dehydrogenase, phosphofructokinase and pyruvate kinase ^12^. In mammalian tissues the synthesis of Glc-1,6-BP is catalyzed by the PGM2L1 enzyme, a member of the cd05799 subfamily, utilizing 1,3-bisphosphoglyceric acid (1,3-BPG) as phosphoryl donor for Glc-1P ^3^. In contrast, despite its essential role in the activity of prokaryotic PGM, the function of Glc-1,6-BP in prokaryotes has been overlooked and its origin remains obscure as no enzyme producing free Glc-1,6-BP has been identified so far ^13^. The unicellular cyanobacterium *Synechocystis* sp. PCC 6803 (hereafter *Synechocystis*) has two enzymes that are encoded by the genes *sll0726* and *slr1334* and annotated as PGM-like ^4^. While Sll0726 exhibits typical properties of PGMs ^14-16^, little is known about Slr1334. The only study addressing this issue reported that purified *Synechocystis* Sll0726 has about a tenfold higher activity *in vitro* than Slr1334 and that a *sll0726* knockout mutant only shows around 3 % PGM activity compared to the wildtype ^17^. Despite its low contribution to over-all PGM activity, the Slr1334 product appeared to be essential as it was not possible to acquire fully segregated knockout mutants of the *slr1334* gene, which raised the question of the role of Slr1334 in *Synechocystis*.

In this work we show that Slr1334 has a phosphotransferase activity catalyzing the production of free Glc-1,6-BP from Glc-1P/Glc-6P and Fructose-1,6-bisphosphate (Frc-1,6-BP). Thereby, Slr1334 is responsible for the formation of the key activator of PGM, Glc-1,6-BP, making it a major regulator in glycolysis and other carbon pathways. Our finding makes Slr1334 the first bacterial enzyme specific for the production of free Glc-1,6-BP. Importantly, we show that the homologue of Slr1334 in the human gut bacterium *Bacteroides salyersiae* catalyzes the same reaction.

## Results

### Classification of the two PGM isoenzymes from *Synechocystis*

*Synechocystis* possesses two genes annotated as PGMs, the genes *sll0726* and *slr1334*. In order to classify them within the αPHM superfamily, we performed a phylogenetic analysis by multi-level alignment of all αPHM subfamilies that are considered PGM or “PGM_like”. For this, we collected the entry sequences from the NCBI Conserved Domains Database (CDD) ^4^ and retrieved further sequences by performing BLAST searches in the NCBI RefSeq_protein database. To focus on glucose-specific αPHMs, the groups of enzymes specific for other sugars like PMMs, PNGM and PAGMs (cd03086 cd03088, cd05802, cd05805) were omitted from the analysis (**Fig. 1**). In general the αPHMs can be divided into four conserved regions which are from N- to C-terminus: (1) the active site domain, (2) the metal binding domain, (3) the sugar binding domain and (4) the phosphate binding domain. While the domains 1,2 and 4 are usually highly conserved between the different subfamilies, the sugar binding domain shows a greater variance and can assist in the categorization of αPHMs (**Table S1**) ^2^.

**Figure 1:**
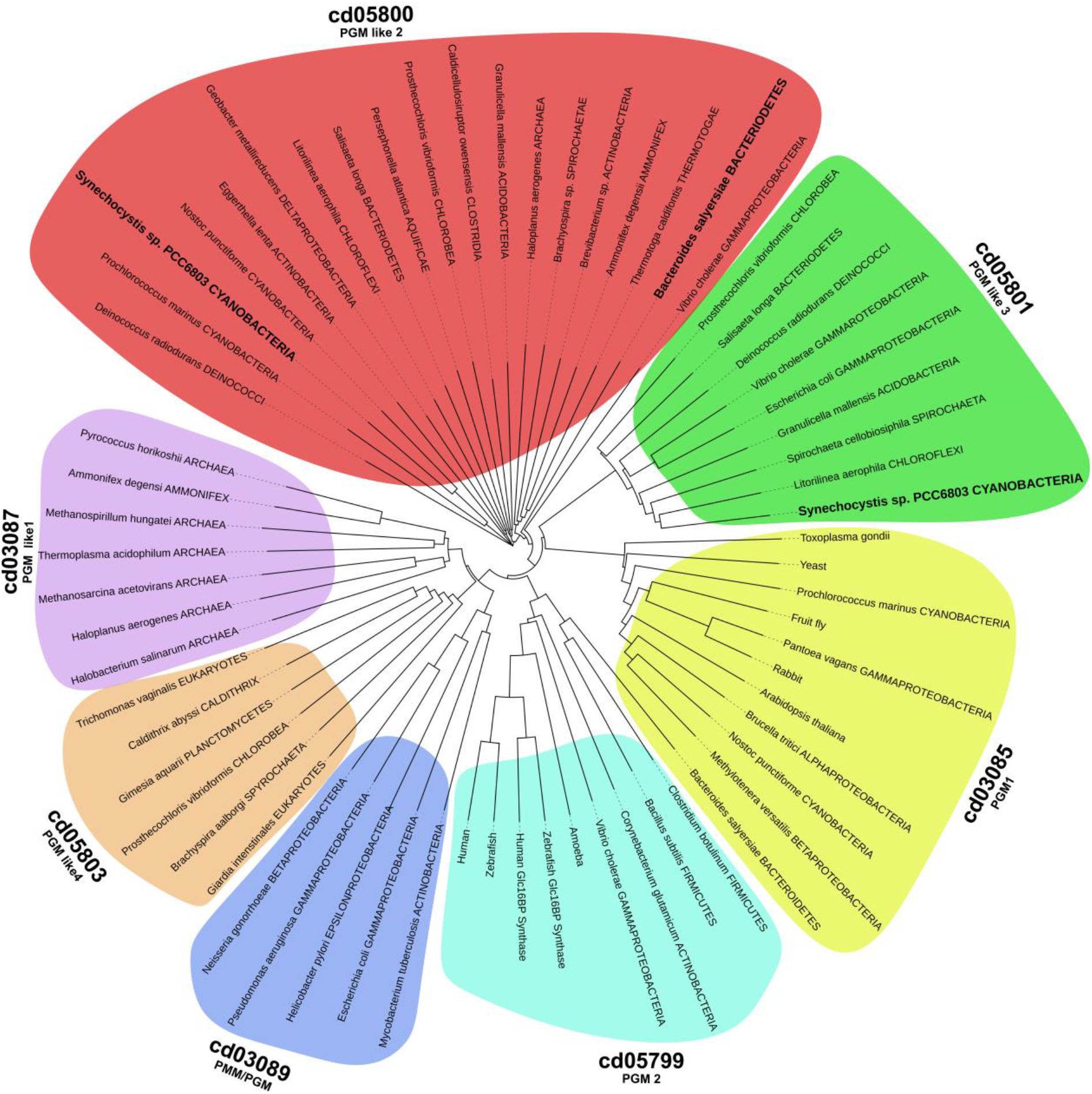
Phylogenetic tree of the αPHM subfamilies that are considered to be “PGM-like” according to CDD. The tree was built on the taxonomy based on the protein sequences present in the NCBI RefSeq_protein database. The CDD subfamilies cd03086, cd03088, cd05802 and cd05805, which are known to catalyze distinct reactions, were omitted from the analysis. The organisms investigated in this study are highlighted in bold letters.

Slr1334 is part of the cd05800 (PGM like 2) subfamily present in many bacterial groups. In agreement with the CDD, our phylogenetic analysis shows that this subfamily forms a distinct cluster of PGMs which clearly separates from other PGM and PGM like enzymes. The Sll0726 PGM, by contrast, is a member of the cd05801 subfamily, to which the well characterized PGM from *S. typhimurium* belongs.

### Slr1334 shows low PGM activity compared to Sll0726 but behaves differently in presence of Fructose-1,6-BP

To gain deeper insights into the role of the enigmatic Slr1334, we overexpressed Sll0726 along with Slr1334 in *Escherichia coli* and assessed the PGM enzymatic activity of the purified proteins by measuring the interconversion of Glc-1-P to Glc-6-P in an assay that couples Glc-6-P formation to its subsequent oxidation using Glucose-6-phosphate-dehydrogenase (G6PDH) and NADP^+^. We could show that Slr1334 only displays about 5% of catalytic efficiency compared to Sll0726 when both PGMs were tested with saturating amounts of the activator Glc-1,6-BP (60 µM) (**Fig. 2, Table 2. See also Fig. 3**). Interestingly, the PGM activity of both enzymes was also measurable without addition of Glc-1,6-BP, which is considered to be an essential activator. However, in the absence of Glc-1,6-BP, the catalytic efficiency was about 70 times lower due to a significantly increased Michaelis-Menten constant (K_m_) for Sll0726 and Slr1334 alike.

**Figure 2:**
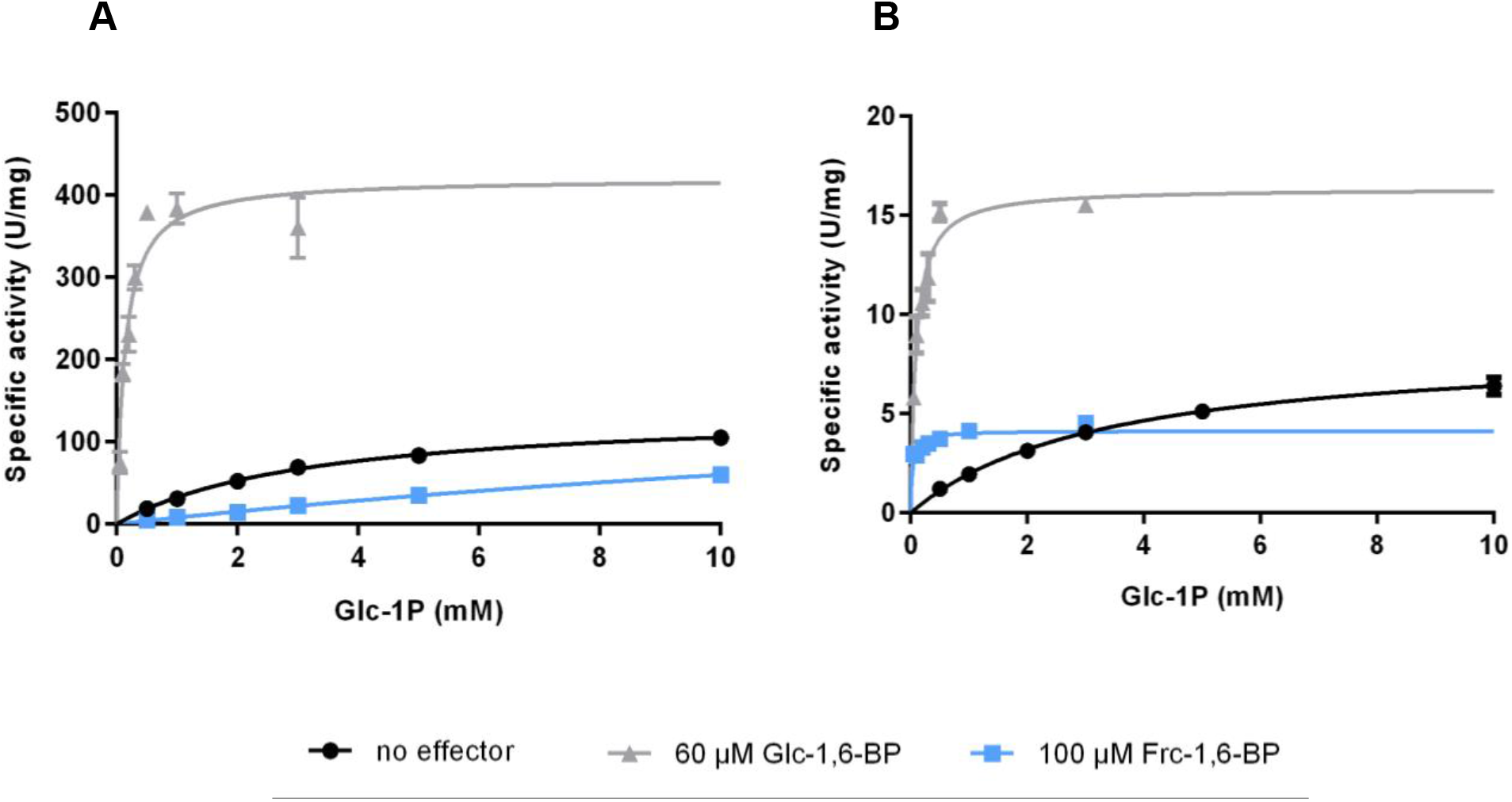
Slr1334 shows lower PGM activity then Sll0726 and behaves differently in presence of Frc-1,6-BP. Shown are the Michaelis-Menten kinetics of Sll0726 (A) and Slr1334 (B): in black, without addition of effectors; in light grey in the presence of 60 µM Glc-1,6-BP; in blue, in the presence of 100 µM Frc-1,6-BP. Three replicates were measured for each data-point. Error bars represent the SD.

**Table 2:** Kinetic parameters of Sll0726 and Slr1334 with different effector molecules. K_m_ and K_cat_ values are means of triplicates with ± SD. Catalytic efficiency ratio is given in percentage and was calculated by the ratio of K_cat_/K_m_ of Slr1334 and Sll0726.

| Effector | SII0726 |  |  | Slr1334 |  |  | Catalytic efficiency ratio (%)<br>(Slr1334/SII0726) |
| --- | --- | --- | --- | --- | --- | --- | --- |
| | $K_m$<br>(mM) | $k_{cat}$<br>($s^{-1}$) | $k_{cat}/K_m$<br>($M^{-1}s^{-1}$) | $K_m$<br>(mM) | $k_{cat}$<br>($s^{-1}$) | $k_{cat}/K_m$<br>($M^{-1}s^{-1}$) | |
| No effector | $3.27 \pm 0.21$ | $138 \pm 3.7$ | $42.2 \cdot 10^3$ | $3.31 \pm 0.14$ | $7.5 \pm 0.1$ | $2.3 \cdot 10^3$ | 5.5 |
| Glc-1,6-BP | $0.19 \pm 0.04$ | $415 \pm 30.6$ | $3038.9 \cdot 10^3$ | $0.09 \pm 0.02$ | $14.4 \pm 0.7$ | $158.1 \cdot 10^3$ | 5.2 |
| Frc-1,6-BP | $26.86 \pm 3.35$ | $218 \pm 21.3$ | $8.1 \cdot 10^3$ | $0.03 \pm 0.01$ | $3.6 \pm 0.2$ | $117.5 \cdot 10^3$ | 1450 |

**Figure 3:**
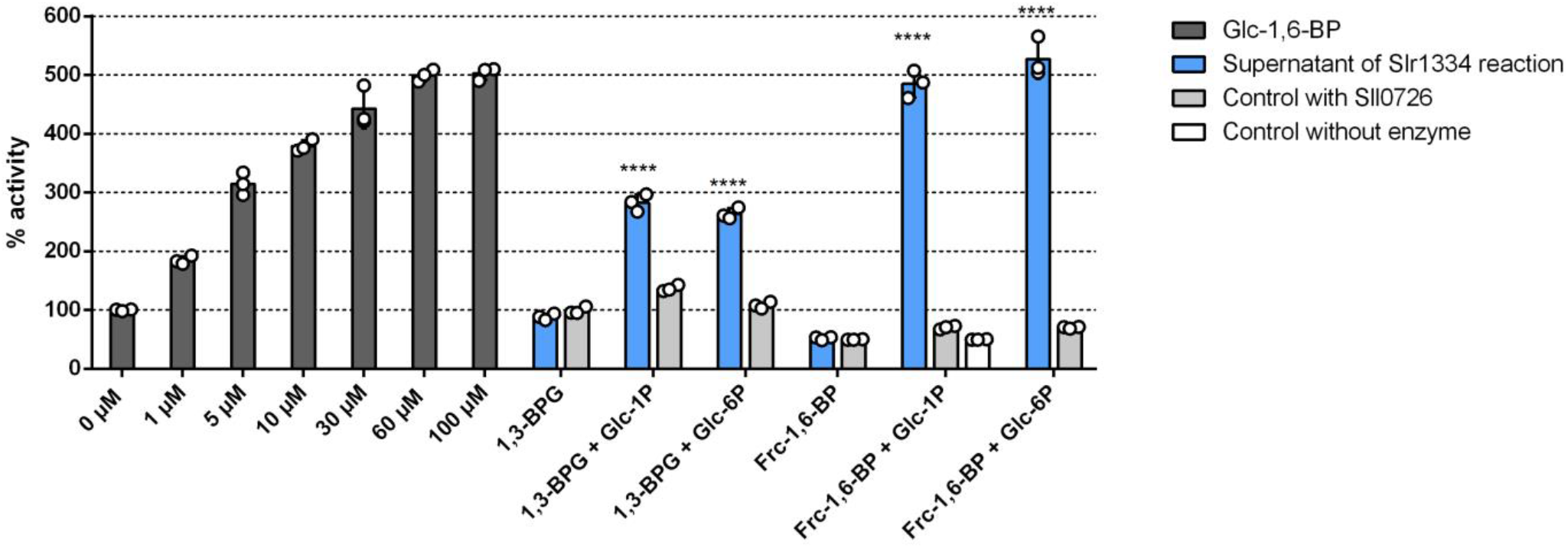
Slr1334 produces a compound out of Frc-1,6-BP that strongly activates Sll0726. Shown is the Sll0726 activity: in dark grey, at different Glc-1,6-BP concentrations; in blue, after addition of the supernatant of the Slr1334 reaction; in light grey, after addition of the supernatant of the Sll0726 reaction as a control; in white, after addition of the sole phosphosugars without enzyme as further control. The different combinations of substrates used for the Slr1334 reaction and controls are indicated on the X-axis. Values represent relative activity compared to the Sll0726 reaction with 0 µM Glc-1,6-BP, which was considered 100%. Three biological replicates were measured for each datapoint. Error bars represent the SD; asterisks represent the statistical significance

Fructose-1,6-bisphosphate is an endogenous intermediate of the glycolytic pathway and is known to be an inhibitor of PGM most likely by competing with the reaction intermediate Glc-1,6-BP ^18, 19^. In a preliminary experiment we tested the effect of different concentrations of Frc-1,6-BP on Sll0726 and observed major inhibition already at concentrations of 100 µM (data not shown). When testing PGM activity by replacing 60 µM Glc-1,6-BP with 100 µM Frc-1,6-BP, we detected a decrease in catalytic efficiency to 19 % as a consequence of a strongly increased K_m_ (**Fig. 2, Table 2**). Strikingly, however, for Slr1334 we detected a different effect: while the catalytic rate constant (k_cat_) decreased by about 2-fold, we measured an approximately 100-fold reduced K_m_ resulting in a 50 times increased catalytic efficiency. This raised the question of how Frc-1,6-BP affected the PGM-activity of Slr1334. We speculated that Frc-1,6-BP might either function as a direct activator similar to Glc-1,6-BP or be required for the synthesis of the activating compound Glc-1,6-BP during the reaction of Slr1334. As we will show below, the activating compound turned out to be Glc-1,6-BP and not Frc-1,6-BP.

### Slr1334 forms a product out of Frc-1,6-BP and Glc-1P/Glc-6P that strongly activates Sll0726

To determine the amount of Glc-1,6-BP needed to achieve activation of Sll0726 at 1 mM Glc-1P we first tested enzymatic activity of Sll0726 at different Glc-1,6-BP concentrations. Maximum activation was achieved at Glc-1,6-BP concentrations between 30 and 60 µM resulting in a 5-fold increased activity as compared to the zero Glc-1,6-BP reaction (**Fig. 3**). Next, we incubated Slr1334 with the phosphorylated sugars Frc-1,6-BP, Glc-1P and Glc-6P (1 mM each) in different combinations and subsequently tested whether Sll0726 could be activated by the corresponding reaction products (**Fig. 3**). As negative control, Frc-1,6-BP and Glc-1P were allowed to react without any enzyme or with Sll0726 instead of Slr1334. After running the Slr1334 reaction for 1.5 h, the reaction was stopped by heat inactivation and the supernatant was added in a dilution of 1:10 to the reaction mixture of the Sll0726 assay, which was then started by adding 1 mM Glc-1P. Only after adding the supernatant of the Slr1334 reaction containing Frc-1,6-BP and Glc-1P (or Glc-6P) as substrates, the velocity of the Sll0726 reaction was increased by 5-fold in comparison to the control reaction at zero Glc-1,6-BP (**Fig. 3**). When the Sll0726 test reaction was performed with the supernatant of the Sll0726 control reaction or the reaction products of Slr1334 using Frc-1,6-BP as sole substrate, the enzyme velocity was lower than the zero Glc-1,6-BP reaction. This reduction can be explained by the inhibitory effect of Frc-1,6-BP on Sll0726, as exemplified by the “no enzyme control” reaction (**Fig. 3**).

In addition to Frc-1,6-BP, we also tested whether Slr1334 can utilize 1,3-BPG instead of Frc-1,6-BP for Glc-1,6-BP formation as described for the mammalian PGM2L1. Since 1,3-BPG is not commercially available, we established an assay whereby we continuously produced 1,3-BPG by a phosphoglycerate kinase (PGK) reaction using a His-tagged purified PGK in combination with 1 mM 3PG and 1 mM ATP (see methods part). This reaction was directly coupled to the reaction of Slr1334 with Glc-1P or Glc-6P or no substrate. As a control the same reaction was performed with Sll0726 instead of Slr1334. To test the possible formation of Glc-1,6-BP, the supernatants of these reactions were added to the Sll0726 reaction before performing the enzymatic assay as described above.

We detected an increase in Sll0726 activity by approx. 2.5-fold compared to the zero Glc-1,6-BP reaction when the supernatant of the Slr1334 reaction contained 1,3-BPG and either Glc-1P or Glc-6P. No increase in activity was detected when the supernatant solely contained 1,3-BPG. The supernatant of the control reaction with Sll0726 instead of Slr1334 resulted in no marked change in activity.

By comparing the activation of Sll0726 by the Slr1334 reaction products to the activation of Sll0726 at different concentration of Glc-1,6-BP we could estimate the efficiency of Glc-1,6-BP formation from Frc-1,6-BP or 1,3-BPG. With Frc-1,6-BP as substrate for the Slr1334 reaction, a maximum activation of Sll0726 was achieved, which corresponded to activation of Sll0726 with concentrations of Glc-1,6-BP between 30 and 60 µM (**Fig. 3**). When 1,3-BPG was used as substrate as described above, the increase in Sll0726 activity corresponded to Glc-1,6-BP concentrations between 1 and 5 µM. From this experiment we conclude that i) Slr1334 forms a product out of Glc-1P/Glc-6P and either Frc-1,6-BP or 1,3-BPG that is able to activate Sll0726 similar to defined amounts of Glc-1,6-BP; ii) when normalizing the Sll0726 activation to Glc-1,6-BP concentrations, the reaction product of Slr1334 using Frc-1,6-BP as phosphorylating sugar is 10-fold more effective in activating Sll0726 than using 1,3-BPG.

### The product of the Slr1334 reaction is not degradable by the Frc-1,6-BP-aldolase

To confirm that the activating compound formed from Glc-1P/Glc-6P and Frc-1,6-BP through the Slr1334 reaction is in fact Glc-1,6-BP, we analyzed the reaction product via liquid chromatography-mass spectrometry (LC/MS). Therefore, we incubated Slr1334 together with 500 µM Frc-1,6-BP and 1 mM Glc-1P for 1.5 h. As a control, the same substrates were incubated without enzyme. Since LC/MS analysis does not allow discriminating Glc-1,6-BP from Frc-1,6-BP, we performed an additional reaction step to specifically degrade Frc-1,6-BP. Therefore, after heat inactivation, the reaction mixture was incubated with Frc-1,6-BP-aldolase (Fba) and glycerol-3-phosphate-dehydrogenase (GPDH) as well as 1 mM NADH: Fba specifically cleaves Frc-1,6-BP into dihydroxyacetone phosphate (DHAP) and glyceraldehyde-3P while Glc-1,6-BP is unaffected. GPDH subsequently forms glycerol-3-phosphate (glycerol-3P) and NAD^+^ out of DHAP and NADH. This is necessary to enable full degradation of Frc-1,6-BP.

After this treatment, LC/MS analysis revealed that a compound with a mass corresponding to either Glc-1,6-BP or Frc-1,6-BP was present in the Slr1334 reaction while this mass was not detectable in the control reaction where no Slr1334 was present (**Fig. 4**). This indicates that Frc-1,6-BP in the control reaction was completely degraded in the subsequent reaction by Fba as expected, while Frc-1,6-BP in the Slr1334 reaction was metabolized into a molecule of the same mass that was not degraded by Fba. This is precisely what is expected when Glc-1,6-BP is produced form Frc-1,6-BP turn-over. Furthermore, the reaction products from the coupled GPDH reaction, NAD+ and glycerol-3P were more abundant in the control reaction without Slr1334 as compared to the reaction with Slr1334. This indicated that more Frc-1,6-BP was degraded out of the control reaction than out of the reaction containing Slr1334 due to partial conversion of Frc-1,6-BP into Glc-1,6-BP by Slr1334 (**Fig. 4**). Altogether, this experiment strongly indicated that in the presence of Slr1334, Frc-1,6-BP was partially utilized to form Glc-1,6-BP. The fact that only a part of Frc-1,6-BP was consumed could be either due to a too short reaction time or because the reaction reached an equilibrium state.

**Figure 4:**
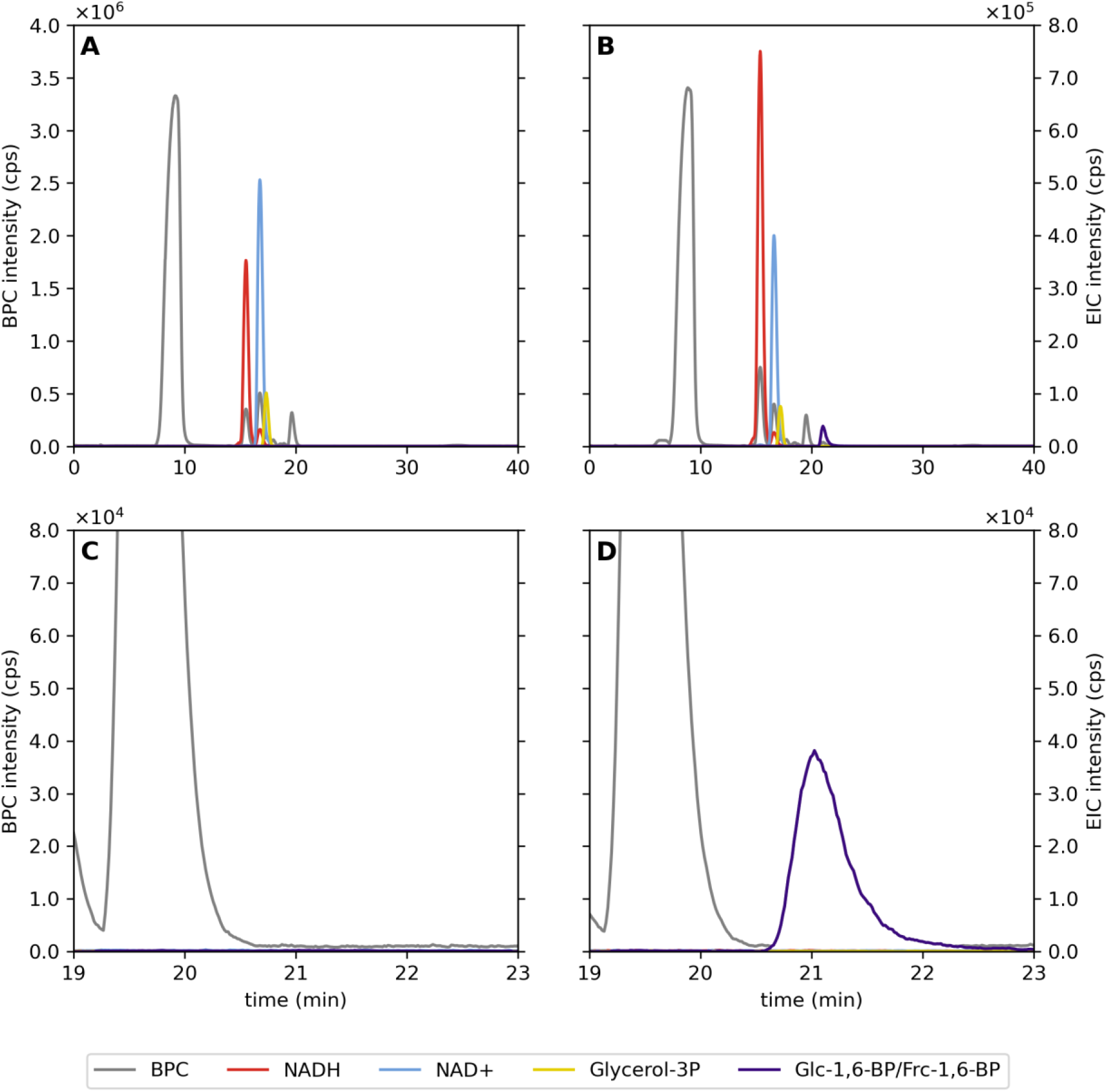
LC-MS analysis reveals that Slr1334 metabolizes Frc-1,6-BP into a molecule of identical mass that is not degraded by Frc-1,6-BP-aldolase. Analysis of the products of the reaction without (A) and with Slr1334 (B) via LC/MS. Shown are the base peak chromatograms (BPC) in the mass range (M-H)^-^ m/z = 79–1,601 (grey) and the extracted ion chromatograms (EIC) based on the exact masses of NAD+ (M-H)^−^= 662.101 m/z (light blue), NADH (M-H)^−^= 664.117 m/z (red), Glycerol 3-P (M-H)^−^= 171.006 m/z (yellow), Glc-1,6-BP/Frc-1,6-BP (M-H)^−^= 338.988 m/z (purple). The presence of the NAD+ (blue) and Glycerol-3P (yellow) peaks in (B) shows that Frc-1,6-BP was not fully metabolized into Glc-1,6-BP. Magnification and adjustment of the graphs in (A) and (B) is clear evidence of the absence (C) or formation (D) of Glc-1,6-BP.

### Slr1334 keeps Frc-1,6-BP and Glc-1,6-BP in an equilibrium

For investigating whether the formation of Glc-1,6-BP out of Glc-1P/Glc-6P and Frc-1,6-BP is an equilibrium reaction, we performed an enzymatic assay to determine the amount of the remaining Frc-1,6-BP. Therefore, Slr1334 was incubated with 500 µM of Frc-1,6-BP and 1 mM Glc-1P for 1.5 h. As controls, the same assays were performed with Sll0726 or without any enzyme. After the reaction was stopped by heat inactivation, the supernatant was used to run the coupled Fba/GPDH assay in a dilution of 1:10, which would correspond to a final concentration of 50 µM Frc-1,6-BP if no Frc-1,6-BP was consumed. As further controls, standards of 50 µM and 25 µM Frc-1,6-BP were used in the Fba/GPDH assay. This experiment showed that the concentration of Frc-1,6-BP decreased by about 50 % when in the first reaction Slr1334 was present, while apparently no Frc-1,6-BP was utilized in the control reactions (**Fig. 5**). This result agreed with the assumption that the reaction reaches an equilibrium at 50 % product formation, either because of the reversibility of the reaction or product inhibition.

**Figure 5:**
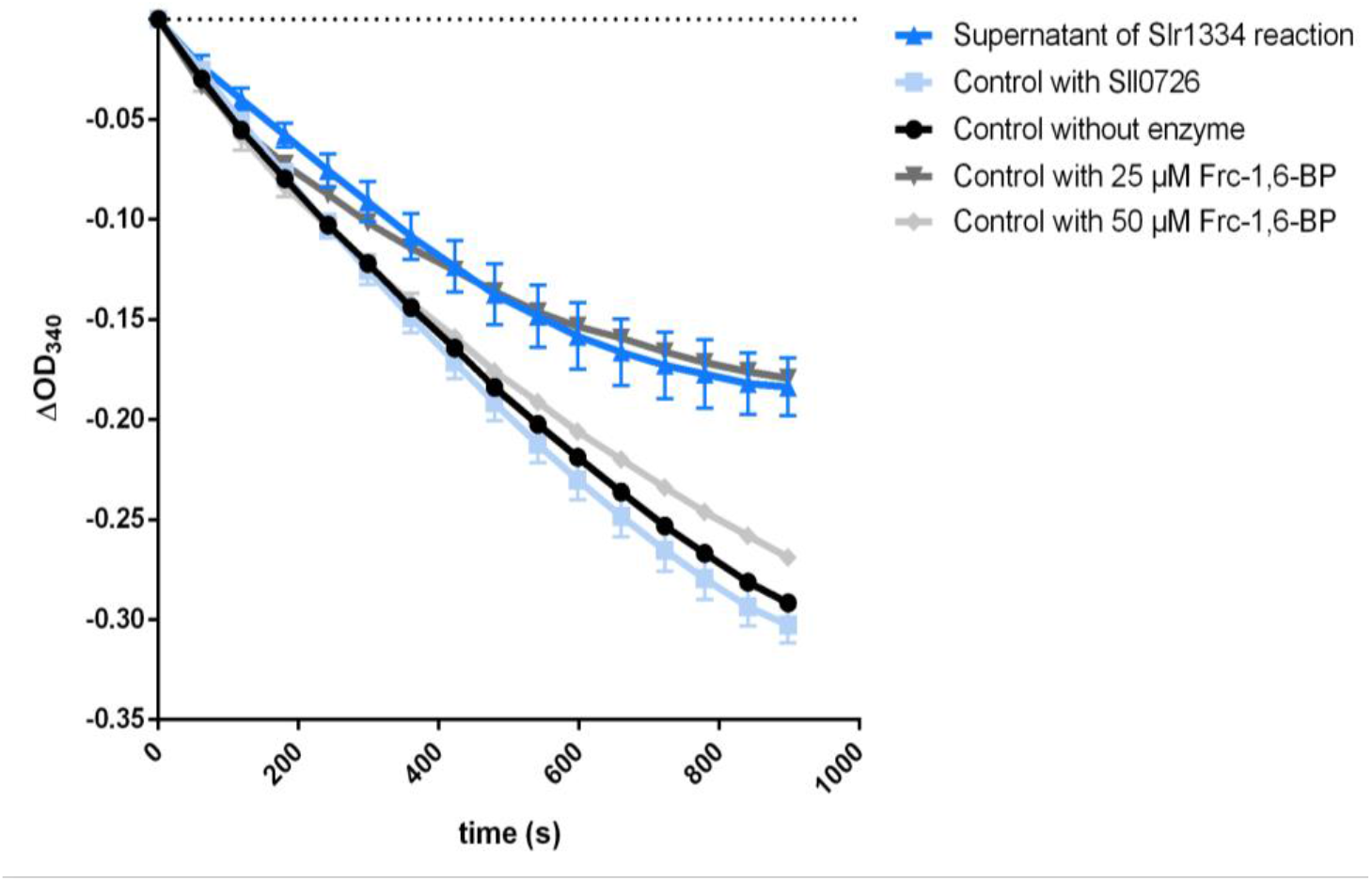
Slr1334 produces Glc-1,6-BP out of Frc-1,6-BP in an equilibrium reaction. Shown is the Frc-1,6-BP-aldolase activity: in dark blue, after addition of the supernatant of the Slr1334 reaction; in light blue, after addition of the supernatant of the Sll0726 reaction as a control; in black, after addition of the supernatant without any enzyme. In dark grey and light grey, the aldolase activity measured at Frc-1,6-BP concentrations of 25 µM and 50 µM, respectively, as further control. Frc-1,6-BP was used at concentrations of 500 µM in the Slr1334 reaction and respective control reactions. Therefore, its final concentration in the Sll0726 assay after 1:10 dilution was expected to be 50 µM assuming no consumption of Frc-1,6-BP in the Slr1334 reaction. Three replicates were measured for each datapoint. Error bars represent the SD.

### The Slr1334 reaction is reversible

To reveal whether Slr1334 can catalyze the proposed reverse reaction and synthesize Frc-1,6-BP from Glc-1,6-BP and Fructose-1P (Frc-1P)/Fructose-6P (Frc-6P), formation of Frc-1,6-BP was coupled to the Fba and GPDH reaction. For the assay, 50 µM Glc-1,6-BP was used and the reaction was started by the addition of 5 mM Frc-6-P. As control reactions, *Synechocystis* Sll0726 as well as PGM1 from rabbit were used. The results clearly showed that Slr1334 could produce Frc-1,6-BP, resulting in measurable aldolase activity while the controls showed no activity (**Fig. 6a**). Using this assay, we tried to determine the kinetic constants for the two substrates Frc-6-P and Glc-1,6-BP. For Frc-6P, a K_m_ of 1 mM and a K_cat_ of 1.2 s^-1^ could be determined (**Fig. 6b, Table 3**). By contrast, the affinity of Slr1334 for Glc-1,6-BP was so high that, even at the lowest measurable substrate concentrations (2 µM), the reaction proceeded with maximal velocity (**Fig. 6c**). Altogether these experiments unequivocally demonstrate that Slr1334 is the first characterized bacterial Glc-1,6-BP synthase. In contrast to the eukaryotic Glc-1,6-BP synthase, the prokaryotic enzyme uses Frc-1,6-BP as major phosphoryl donor for Glc-1,6-BP production.

**Figure 6:**
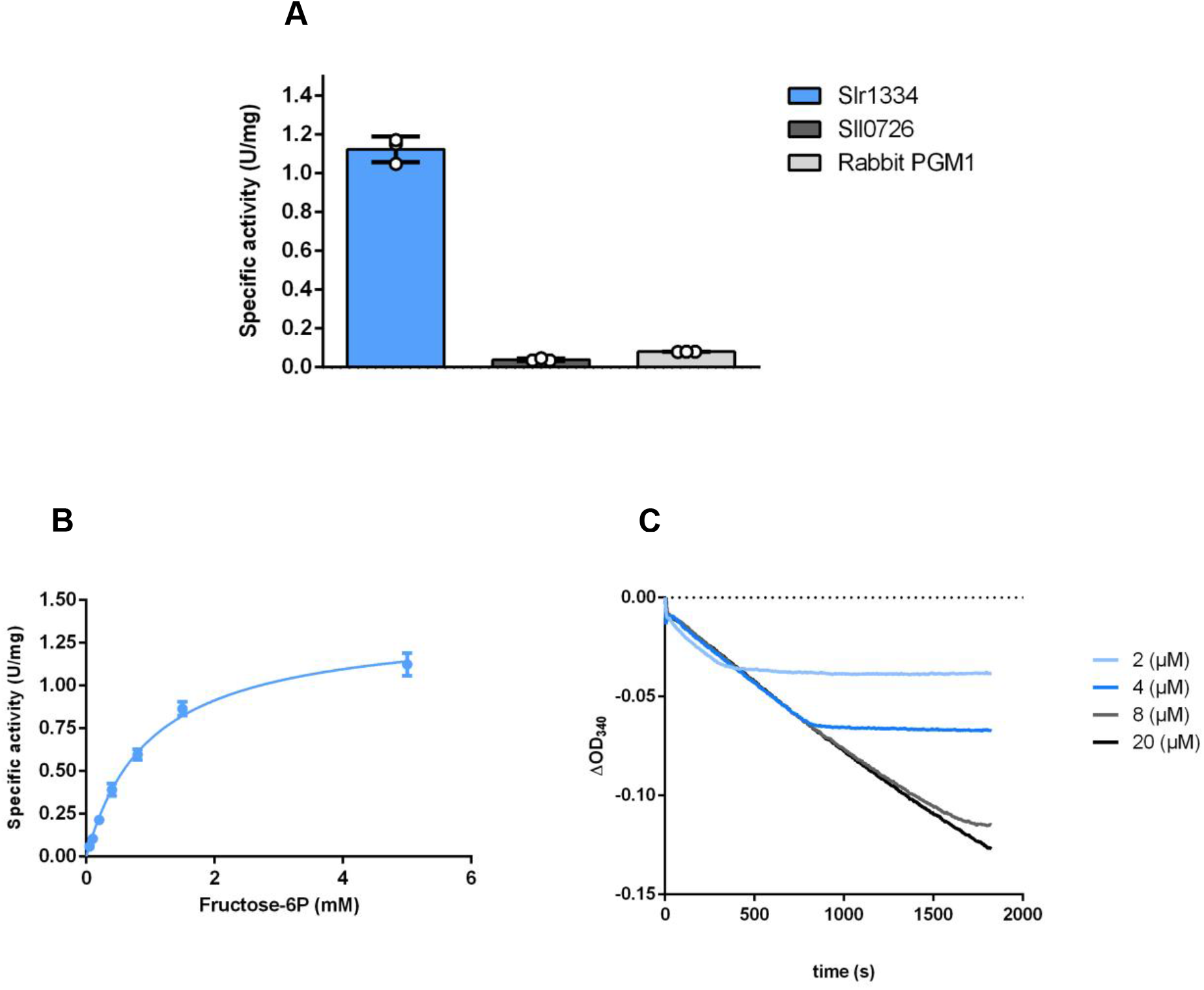
Slr1334 catalyzes the reverse reaction to form Frc-1,6-BP out of Glc-1,6-BP and Frc-6P. (A) Activity of Slr1334 (blue), Sll0726 (grey), and rabbit PGM1 (black) using 50 µM Glc-1,6-BP and 5 mM Frc-6P. (B) Michaelis-Menten kinetics of Slr1334 using different concentrations of Frc-6P and 50 µM Glc-1,6-BP. (C) Slr1334 reaction at different concentrations of Glc-1,6-BP and 0.5 mM Frc-6P. Three replicates were measured for each datapoint. Error bars represent the SD; asterisks represent the statistical significance.

**Table 3:** Kinetic parameters of Frc-1,6-BP synthesis by Slr1334. K_m_ and Kcat values are means of triplicates with ± SD.

| Substrate | $K_m$ (mM) | $k_{cat}$ ($s^{-1}$) | $k_{cat}/K_m$ ($M^{-1}s^{-1}$) |
| --- | --- | --- | --- |
| Glc-1,6-BP | <0.002 | $1.2 \pm 0.03$ | 600000 |
| Frc-6P | $1.0 \pm 0.07$ | | >1200 |

### The Glc-1,6-BP synthase reaction is also performed by other members of the cd05800 group

To determine whether the reaction catalyzed by Slr1334 is representative for αPHM enzymes of the cd05800 family, we decided to analyze the corresponding homologue from the human microbiome associated bacterium *Bacteroides salyersiae*. Therefore, we tested formation of Glc-1,6-BP under the same conditions as described above for Slr1334 by incubating the recombinant cd05800 enzyme from *B. salyersiae* with Frc-1,6-BP and Glc-1P or Glc-6P and subsequently adding the supernatant of this reaction to the Sll0726 assay. As a positive control we used the reaction of Slr1334.

The addition of the supernatant from the *B. salyersiae* cd05800 PGM increased the velocity of Sll0726 in a comparable manner as the supernatant of the Slr1334 reaction (**Fig. 7**). This indicates that similar amounts of Glc-1,6-BP have been formed in both reactions, hence revealing that *B. salyersiae* cd05800 PGM catalyzes the same reaction as Slr1334.

**Figure 7:**
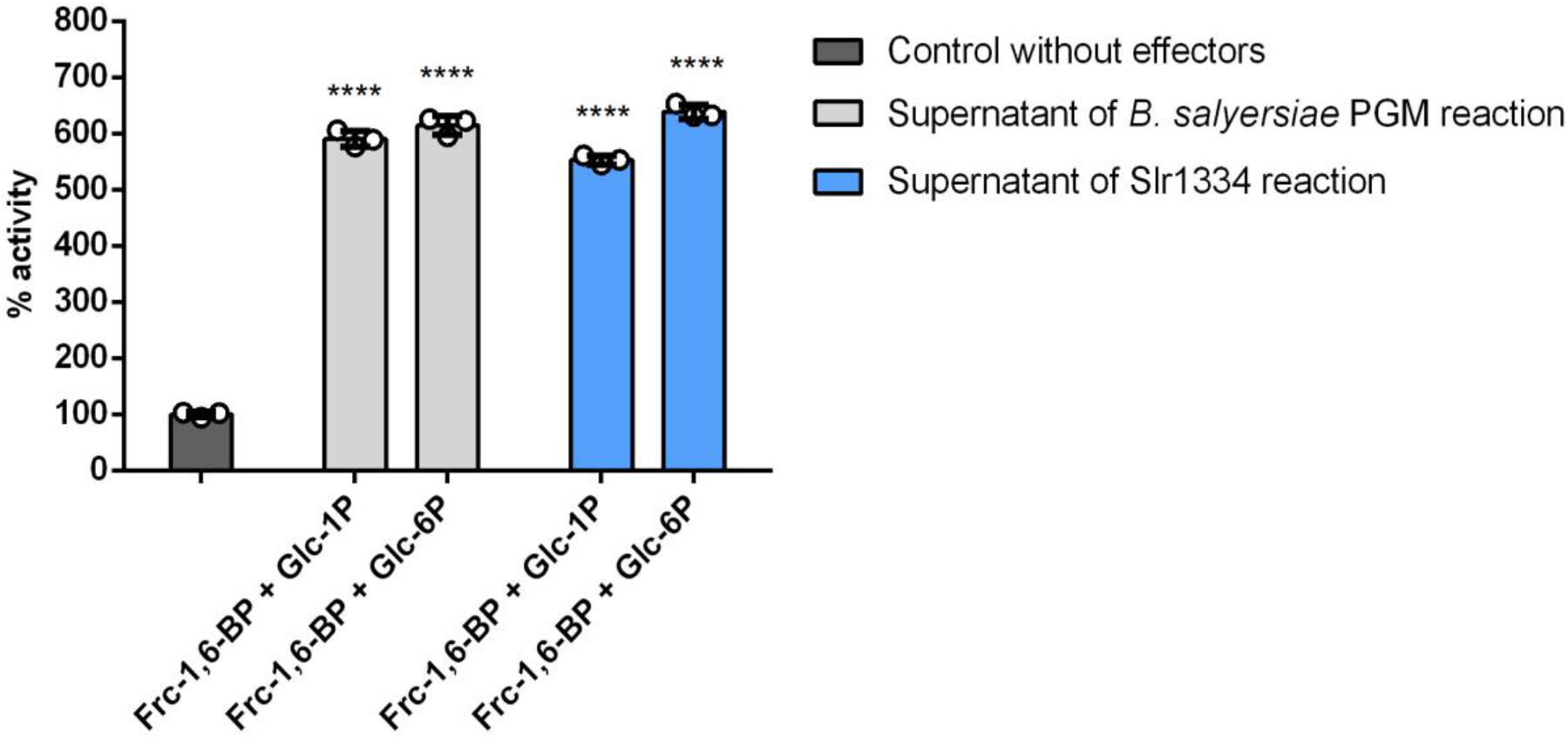
Slr1334 homologue from *B*.*salyersiae* also forms Glc-1,6-BP out of Frc-1,6-BP. Shown is the activity of Sll0726: blue, after addition of the supernatant of the *B. salyersiae* PGM reaction; in light grey, after addition of the supernatant of the Slr1334 reaction as a positive control; in dark grey, without any effectors as a negative control. The different combinations of substrates used for the *B. salyersiae* PGM and Slr1334 control reaction are indicated on the X-axis. Values represent relative activity compared to the Sll0726 reaction without any effectors, which was considered 100%. Error bars represent the SD; asterisks represent the statistical significance.

## Discussion

Among the key reactions in carbon metabolism are the phosphoryl transfer reactions on different phosphosugars, which are catalyzed by the members of the αPHM superfamily. These reactions are essential for glycogen metabolism, amino sugar metabolism as well as cell wall synthesis and are conserved throughout all domains of life. The interconversion of Glc-1P to Glc-6P and vice versa by PGM is by far the best studied of these reactions and investigation of PGM goes back to the 1940s. It has long been known that PGM requires Glc-1,6-BP as a activator ^10^. While in mammals free Glc-1,6-BP has been shown to derive from the reaction of the PGM2L1 enzyme utilizing Glc-1P and 1,3-BPG as substrates, its origin in prokaryotes has remained unknown ^3^.

In this study we used the unicellular cyanobacterium *Synechocystis* sp. PCC6803 as a model organism to identify the first bacterial glucose-1,6-BP synthase. This reversible reaction is a novel function of a member of a distinct αPHM subfamily producing free α-D-Glc-1,6-BP and Frc-6P from Frc-1,6-BP and Glc-1P/Glc-6P.

Glc-1,6-BP is essential for PGM activity due to the requirement for phosphorylation of the serine residue at the active site for enzyme activation. The activating reaction converts Glc-1,6-BP into either Glc-1P or Glc-6P. After this initial activation, the interconversion of Glc-1P to Glc-6P is mediated by a transfer of the phosphate group from the serine residue at the active site to the 1’ or 6’ position of the substrate respectively, forming a Glc-1,6-BP intermediate. This intermediate has to reconfigure inside the enzyme and transfer the phosphate of its 6’ or 1’ position back to the active site serine leading to the release of Glc-6P or Glc-1P, respectively. As a result of this mechanism, the active site remains phosphorylated and the enzyme is ready for a new cycle. However, it has been shown that the phosphorylation of PGM at the active site is lost after an average of 15-20 reaction cycles by a spontaneous dissociation of the intermediate Glc-1,6-BP leaving behind an inactive dephosphoenzyme. Therefore, a constant level of Glc-1,6-BP is necessary to ensure that PGM is kept in an active state ^20, 21^. While the loss of the intermediate might be regarded as an unfavorable event, it can also be considered as a regulatory mechanism that enables regulation of PGM activity based on Glc-1,6-BP levels. For other αPHMs like PMM and PNGM it is known that the corresponding intermediates (e.g. α-D-mannose-1,6-BP and α-D-glucosamine-1,6-BP) are formed during the reaction of the respective enzymes in the same way as for PGM but no source of those intermediates is known that would enable an initial activation of these αPHMs or would keep them in an active state overtime when phosphorylation is lost. However, since many members of the αPHM family show at least some residual PGM activity, Glc-1,6-BP was suggested to be the activating compound for these enzymes as well. This has in fact been shown for PNGM from *E. coli* which is considered to be an essential enzyme ^22^. This makes the role of Glc-1,6-BP in carbon metabolism even more relevant and would explain the failed attempts to delete the *slr1334* gene ^17^.

In mammals, it was shown that Glc-1,6-BP can also regulate central enzymes of carbon pathways by inhibiting hexokinases, 6-phosphogluconate dehydrogenase and fructose-1,6-bisphosphatase and by activating phosphofructokinase and pyruvate kinase ^12^. Further studies in mammals showed that Glc-1,6-BP is especially elevated in brain tissue to a level that highly exceeds the regulatory concentrations needed for PGM activity, and that reduced levels of Glc-1,6-BP cause severe neurological problems ^23^. This indicates that at least in mammals there are likely more functions of Glc-1,6-BP to be discovered. Given the important role of Glc-1,6-BP in eukaryotic metabolism, it is surprising that very little is known on the role of Glc-1,6-BP in bacterial physiology. The only study about the origin of α-D-Glc-1,6-BP production in bacteria proposed a phosphodismutase reaction with Glc-1P as the sole substrate in *E. coli* ^*24*^. Moreover, the levels of Glc-1,6-BP in eukaryotes are affected by a specific Glc-1,6-BP phosphatase, PMM1, which is a phosphomannomutase that is not part of the αPHM superfamily. Whether a homologue exists in bacteria is currently unknown ^25^.

In absence of Glc-1,6-BP, both Slr1334 and Sll0726 showed low PGM activity. This residual activity can be explained by the fact that a phosphorylation of the active site serine can also be achieved at high concentrations of Glc-1P but at a much lower efficiency, leading to high K_m_ values and overall strongly reduced activity ^11^. In the presence of Frc-1,6-BP, Sll0726 was strongly inhibited, which agrees with the inhibitory effect of Frc-1,6-BP on other PGMs ^19, 26^. By contrast, Frc-1,6-BP enhanced the overall catalytic activity of Slr1334, primarily because of a strongly decreased K_m_. This effect can now be attributed to the formation of the activator Glc-1,6-BP.

Like mammalian PGM2L1, Slr1334 can utilize 1,3-BPG as a phosphorylating compound for Glc-1,6-BP formation, however with a much lower efficiency than with Frc-1,6-BP. For the mammalian PGM2L1 a reverse reaction (formation of 1,3-BPG and Glc-1P from Glc-1,6-BP and 3-PG) has not been shown. For Slr1334, we could clearly show reversibility of the reaction, producing Frc-1,6-BP from Glc-1,6-BP and Frc-6P. Provided that both directions of the reaction have similar kinetic properties, Slr1334 appears to be a *bona-fide* Glc-1,6-BP synthase, which is prone to activate the main PGM Sll0726.

In contrast to Glc-1,6-BP, Frc-1,6-BP is an obligatory intermediate in glycolysis, gluconeogenesis and Calvin-Benson-Bassham cycle. In contrast to mammalian PGM2L1, *Synechocystis* Slr1334 uses this sugar as the major substrate for Glc-1,6-BP synthesis. This links the synthesis of Glc-1,6-BP to the levels of Frc-1,6-BP. Its steady state level depends on the activity of the opposing reactions of phosphofructokinase (Pfk) and Fructose-1,6-bisphosphatase (FBPase), both of which are regulated in an opposite manner: While PFK is usually activated by the low energy state metabolites ADP and AMP and downregulated by the high energy state metabolites ATP and citrate, FBPase is downregulated by AMP while unaffected by high energy compounds. The level of Frc-1,6-BP is further determined by the Frc-1,6-BP-aldolase activity. Frc-1,6-BP is also known to be an allosteric activator of pyruvate kinase (PK), the final step in glycolysis ^27^.

Altogether, in the cyanobacterium *Synechocystis* PCC6803, Frc-1,6-BP appears to have two counteracting effects on PGM activity. On the one hand, it acts as a direct inhibitor of PGM; on the other hand, it gives rise to the synthesis of the PGM activator Glc-1,6-BP through the activity of Slr1334. Furthermore, since these double phosphorylated sugars may affect further enzyme reactions, by connecting these two metabolites, Slr1334 might represent a crucial regulatory point in glycolysis (**Fig. 8**). This conclusion would further underline the essential role of *slr1334* ^17^.

**Figure 8:**
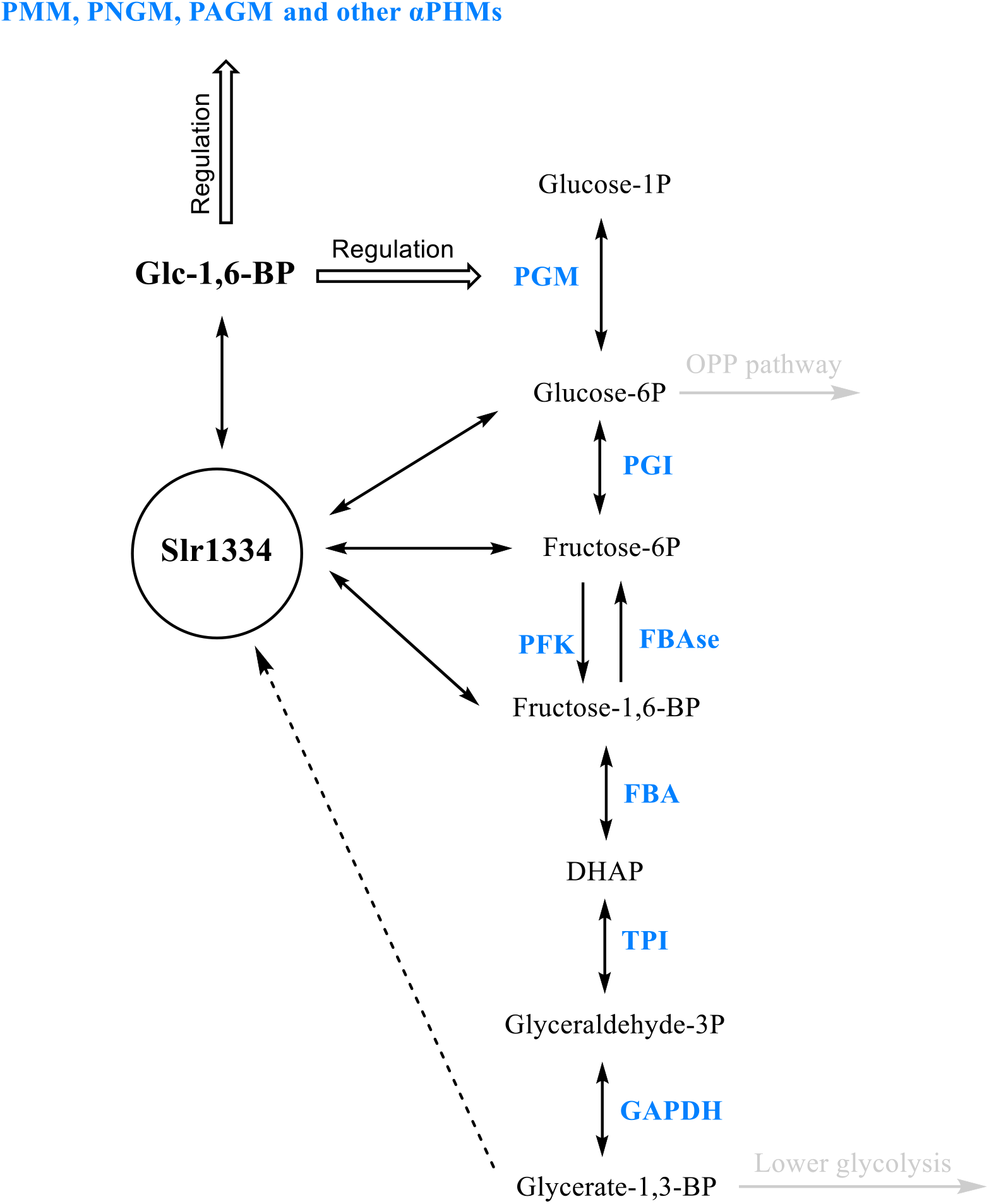
Potential role of Slr1334 in glycolysis. Shown is the potential role of Slr1334 in regulating activity of αPHMs and the concentration of key metabolites in glycolysis. PGM: Phosphoglucomutase; PGI: Glc-6P-Isomerase; PFK: Phosphofruktokinase; FBAse: Fructose-6P phosphatase; FBA: Frc-1,6-BP-aldolase; TPI: Triosephosphate isomerase; GAPDH: Glyceraldehyde-3P dehydrogenase; DHAP: Dihydroxyacetone phosphate

Interestingly, our phylogenetic analysis revealed the cd05800 subfamily as the most primordial group of phosphoglucomutases. This group appears to be especially present in bacterial classes that are considered deeply branching bacteria meaning they are relatively close to the last common universal ancestor (LUCA). Examples for this are the classes of Deinococci, Aquificae, Thermotogae, Bacteroidetes and Cyanobacteria, where this PGM subfamily is widely distributed. This finding is supported by the fact that cd05800 members can also be found in archaea. In agreement with this, the first characterized representative of this group, Slr1334 shows both PGM and Glc-1,6-BP activity. This suggests that in primordial systems these PGMs with broad functionality could have performed PGM reactions without the requirement for specific Glc-1,6-BP synthase enzymes. During subsequent functional diversification, the PGMs may have adopted unique functions, with specialization towards efficient conversion of Glc-6-P into Glc-1P on the one hand, and more specialized formation of the activating compound Glc-1,6-BP on the other hand. This assumption agrees with the observation that many bacteria that have a cd05800 family member also possess a specialized PGM of the cd05801 or cd03085 subfamily for efficient Glc-6P – Glc-1P conversion. This includes the ensemble of all cyanobacteria. Conversely, many pathogenic bacteria that have a PGM of the cd03089 type lack a cd5800 representative (**Table S4**) which raises the question how those strains produce Glc-1,6-BP. To clarify these issues further investigations on αPHM members from various bacterial groups are required.

We were able to show that the cd05800 PGM from the human gut commensal bacterium *B. salyersiae* also catalyzes the synthesis of Glc-1,6-BP out of Frc-1,6-BP like Slr1334 (**Fig. 7**). In the case of *B. salyersiae*, the enzyme appears to have an even higher catalytic efficiency for the Glc-1,6-BP synthase reaction than for the standard PGM reaction (**Fig. S1**). Therefore, we conclude that the formation of Glc-1,6-BP is a general feature of this group of PGMs.

Interestingly, members of the cd05800 group are also present in emerging human pathogenic species like *Vibrio, Eggerthella* and *Brachyospyra* (**Fig. 1**) and are likely to play a key role in the metabolism of these pathogens. In fact, due to their crucial role in capsule and cell wall synthesis PGMs have been suggested as novel targets for the development of antiinfectives in prophylactic and therapeutic interventions ^5, 28^. It is tempting to speculate that Glc-1,6-BP synthases of the cd05800 family, in virtue of their absence in eukaryotes, may also be interesting candidates for the development of novel drugs modulating the microbiota with little collateral damage to the host.

## Methods

### Isothermal, Single-Reaction DNA Assembly (Gibson Cloning)

Cloning was performed as described by Gibson, Young ^29^ using *E. coli* NEB10β cells (details in **Table S5**). All primers and plasmids used are shown in **Table S3** and **Table S4**, respectively.

### Cultivation of *Escherichia coli*

If not otherwise stated *E. coli* was grown in Luria-Bertani medium at 37°C ^30^. For growth on plates, 1.5% (w/v) agar-agar was added. For cells containing plasmids, the appropriate concentration of antibiotics was used. All *E. coli* strains used in this study are listed in **Table S5**.

### Protein overexpression and purification

The plasmids used for protein overexpression are shown in **Table S4**. *Escherichia coli* Rosetta-gami (DE3) (details **Table S5**) was used for the overexpression of all proteins. For this, cells were cultivated in 2xYT (3.5% tryptone, 2% yeast extract, 0.5% NaCl; 1L of culture in 5L flasks) at 37 °C until reaching exponential growth (OD_600_ 0.6-0.8). Protein overexpression was induced by adding 75 µg/L anhydrotetracycline, followed by incubation at 20°C for 16 h. Cells were harvested by centrifugation at 4000 g for 10 min at 4 °C. Cell disruption was performed by sonication in 40 mL of lysis buffer (100 mM Tris-HCl pH 7.5, 500 mM NaCl, 10 mM MgCl_2,_ 20 mM Imidazole for His-tagged proteins and 100 mM Tris-HCl pH 8, 150 mM NaCl, 10 mM MgCl_2_ for Strep-tagged proteins) Lysis buffers were supplemented with DNAse I, and cOmplete^(tm)^ protease inhibitor cocktail (Roche, Basel)). The cell lysate was centrifuged at 40,000 g for 45 min at 4°C and the supernatant was filtered with a 0.22 µM filter.

For the purification of His-tagged proteins, 5 mL Ni-NTA HisTrap columns (GE Healthcare, Illinois, USA) were used. The cell extracts were loaded onto the columns, washed with wash buffer (50 mM Tris-HCl pH 7.4, 500 mM NaCl, and 50 mM Imidazole) and eluted with elution buffer (50 mM Tris-HCl pH 7.4, 500 mm NaCl, and 500 mM Imidazole).

### Measurement of PGM activity of Sll0726 and Slr1334 in vitro

Buffer for enzymatic reactions was composed of 50 mM HEPES-KOH pH 7.5, 150 mM KCl, 10 mM MgCl_2_, 1 mM NADP^+^, 1 mM DTT and 1 U/mL G6PDH from *Saccharomyces cerevisiae* (G6378, Sigma Aldrich). For Sll0726 activity, 60 ng of Strep-tagged purified protein was added to each reaction. For slr1334 activity, 650 ng of purified protein was added. For tests with saturating Glc-1,6-BP concentrations, 60 µM Glc-1,6-BP have been used. Reaction was started by the addition of glucose-1P. Reactions were carried out in a total of 300 µl in a 96-well microplate. Absorption change at 340 nm was continuously measured for 15 min at 30 °C in a TECAN Spark® Multiplate reader (Tecan Group AG, Männedorf, Switzerland). At least three replicates were measured.

### Measurement of Glc-1,6-BP formation

The reaction buffer was composed of 50 mM HEPES-KOH pH 7.5, 150 mM KCl and 10 mM MgCl_2_. To test the formation of Glc-1,6-BP out of Frc-1,6-BP 1 mM Frc-1,6-BP, 1 mM Glc1P or 1 mM Glc-6P were added in different combinations. To test the formation of Glc-1,6-BP out of 1,3-BPG instead of Frc-1,6-BP, 1 mM of ATP, 1 mM 3-PG and 60 µg PGK have been used. Sll0726 and Slr1334 concentration were 30 µg/ml. The reactions were performed for 1.5 h at 30 °C followed by heat inactivation at 90°C for 10 minutes with subsequent centrifugation at 25000 g for 10 minutes at 4° C. The supernatants of the various reactions were used at 1:10 dilution in the Sll0726 activity assay.

### Frc-1,6-BP aldolase assay

To measure the formation of Frc-1,6-BP by Slr1334, the reaction buffer was composed of 50 mM HEPES-KOH pH 7.5, 150 mM KCl, 10 mM MgCl_2_, 0.2 mM NAD, 7 µg Slr1334, 1U/ml aldolase from rabbit muscle and 1U/ml GDH from rabbit muscle. Concentrations of Glc-1,6-BP and Frc-6P were varying depending on the experiment. Reaction was started by the addition of Glc-1,6-BP or Frc-6P depending on the experiment.

To measure residual Frc-1,6-BP after running the Slr1334 reaction, the buffer was composed of 50 mM HEPES-KOH pH 7.5, 150 mM KCl, 10 mM MgCl_2_, 0.2 mM NAD, 0.02U/ml aldolase from rabbit muscle and 1U/ml GDH from rabbit muscle.

All reactions were carried out in a total of 300 µl in a 96-well microplate Absorption change at 340 nm was continuously measured for 15 min at 30 °C in a TECAN Spark® Multiplate reader (Tecan Group AG, Männedorf, Switzerland). The enzymatic activity was then calculated. At least three replicates were measured.

### LC-MS measurement

#### Sample preparation

Reaction buffer was composed of 50 mM HEPES-KOH pH 7.5, 150 mM KCl and 10 mM MgCl_2_. For the reaction without enzyme 500 µM Frc-1,6-BP and 1 mM Glc1-P were added. For the Slr1334 reaction 30 µg of Slr1334 have been used additionally. Reaction was carried out in a total volume of 1 ml at 30 °C for 1h. Afterwards reaction was stopped by heat inactivation at 90° C followed by centrifugation at 25000 g for 10 minutes at 4°C.

For degradation of Frc-1,6-BP 1 mM NADH, 2U/ml of aldolase from rabbit muscle and 2U/ml of GDH from rabbit muscle were added. Reaction was carried out followed by inactivation at the same conditions as before.

#### LC/MS analysis

LC/MS analysis was performed using an electrospray ionization time of flight (ESI-TOF) mass spectrometer (MicrOTOF II; Bruker Daltonics), operated in negative ion-mode connected to an UltiMate 3000 high-performance liquid chromatography (HPLC) system (Dionex). The separation in the HPLC was carried out using a SeQuant ZIC-pHILIC column (PEEK 150 × 2.1 mm, 5 μm, 110 Å, Merck) at 30°C with an CH_3_CN (buffer A) and 100 mM (NH_4_)_2_CO_3_, pH 9 (buffer B) buffer system. A single run (injection volume of 5 μl) was performed with a flow rate of 0.2 ml/min and a linear gradient of 25 min, reducing the concentration of buffer A from 82 to 42 %. Before (5 min) and after (10 min) the gradient, the column was equilibrated with 82 % buffer A.

### Phylogenetic analysis of αPHM subfamilies

The occurrence of members of the αPHM subfamilies was investigated in organisms included in the NCBI RefSeq_protein database. Phylogenetic tree data were created using the ClustalW implementation from the European Bioinformatics Institute (EMBL-EBI) and the phylogenetic tree was created using the Interactive Tree Of Life (iTOL) v5 online tool ^31, 32^.

### Statistical analysis

Statistical details for each experiment can be found in the figure legends. GraphPad PRISM was used to perform one-sided ANOVA to determine the statistical significance. Asterisks (*) in the figures symbolize the p-value: One asterisk represents p ≤ 0.05, two asterisks p ≤ 0.01, three asterisks p ≤ 0.001, and four asterisks p ≤ 0.0001

## Acknowledgements

We thank Libera Lo Presti for her assistance in writing this manuscript and Gaia Bianchi for support in performing enzymatic assays.

This work was supported by FOR2816 “The Autotrophy-Heterotrophy Switch in Cyanobacteria: Coherent Decision-Making at Multiple Regulatory Layers” and EXC2121 “Controlling Microbes to Fight Infections (CMFI).

## Author contributions

N.N. performed cloning, protein purification, enzymatic assays phylogenetic analysis and wrote manuscript. S.F. performed LC/MS analysis. K.F. conceived study, interpreted data and edited manuscript.

## References

1. Shackelford, G.S., C.A. Regni, and L.J. Beamer, Evolutionary trace analysis of the α-D-phosphohexomutase superfamily. Protein Science, 2004. 13(8): p. 2130–2138.

2. Stiers, K.M., A.G. Muenks, and L.J. Beamer, Biology, Mechanism, and Structure of Enzymes in the α-d-Phosphohexomutase Superfamily. Advances in protein chemistry and structural biology, 2017. 109: p. 265–304.

3. Maliekal, P., et al., Molecular identification of mammalian phosphopentomutase and glucose-1,6-bisphosphate synthase, two members of the alpha-D-phosphohexomutase family. J Biol Chem, 2007. 282(44): p. 31844–51.

4. Lu, S., et al., CDD/SPARCLE: the conserved domain database in 2020. Nucleic Acids Res, 2020. 48(D1): p. D265–d268.

5. Regni, C., P.A. Tipton, and L.J. Beamer, Crystal structure of PMM/PGM: an enzyme in the biosynthetic pathway of P. aeruginosa virulence factors. Structure, 2002. 10(2): p. 269–79.

6. Regni, C., et al., Structural basis of diverse substrate recognition by the enzyme PMM/PGM from P. aeruginosa. Structure, 2004. 12(1): p. 55–63.

7. Akutsu, J.-i., et al., Characterization of a Thermostable Enzyme with Phosphomannomutase/Phosphoglucomutase Activities from the Hyperthermophilic Archaeon Pyrococcus horikoshii OT3. The Journal of Biochemistry, 2005. 138(2): p. 159–166.

8. Liu, Y., W.J. Ray, Jr., and S. Baranidharan, Structure of rabbit muscle phosphoglucomutase refined at 2.4 A resolution. Acta Crystallogr D Biol Crystallogr, 1997. 53(Pt 4): p. 392–405.

9. Backe, P.H., et al., Structural basis for substrate and product recognition in human phosphoglucomutase-1 (PGM1) isoform 2, a member of the α-d-phosphohexomutase superfamily. Scientific Reports, 2020. 10(1): p. 5656.

10. Leloir, L.F., R.E. Trucco, and et al., The co-enzyme of phosphoglucomutase. Arch Biochem, 1948. 19(2): p. 339.

11. Naught, L.E. and P.A. Tipton, Kinetic Mechanism and pH Dependence of the Kinetic Parameters of Pseudomonas aeruginosa Phosphomannomutase/Phosphoglucomutase. Archives of Biochemistry and Biophysics, 2001. 396(1): p. 111–118.

12. Carreras, J., et al., Bisphosphorylated metabolites of glycerate, glucose, and fructose: Functions, metabolism and molecular pathology. Clinical Biochemistry, 1986. 19(6): p. 348–358.

13. Joshi, J.G. and P. Handler, PHOSPHOGLUCOMUTASE. I. PURIFICATION AND PROPERTIES OF PHOSPHOGLUCOMUTASE FROM ESCHERICHIA COLI. J Biol Chem, 1964. 239: p. 2741–51.

14. Lindahl, M. and F.J. Florencio, Thioredoxin-linked processes in cyanobacteria are as numerous as in chloroplasts, but targets are different. Proceedings of the National Academy of Sciences, 2003. 100(26): p. 16107–16112.

15. Mehra-Chaudhary, R., et al., Crystal structure of a bacterial phosphoglucomutase, an enzyme involved in the virulence of multiple human pathogens. Proteins, 2011. 79(4): p. 1215–1229.

16. Doello, S., et al., Regulatory phosphorylation site tunes Phosphoglucomutase 1 as a metabolic valve to control mobilization of glycogen stores. bioRxiv, 2021: p. 2021.04.15.439997.

17. Lian Liu, H.H., Hong Gao, and Xudong Xu, Role of two phosphohexomutase genes in glycogen synthesis in Synechocystis sp. PCC6803. Chinese Science Bulletin, 2013. 58(36).

18. Maino, V.C. and F.E. Young, Regulation of Glucosylation of Teichoic Acid: II. PARTIAL CHARACTERIZATION OF PHOSPHOGLUCOMUTASE IN BACILLUS SUBTILIS 168. Journal of Biological Chemistry, 1974. 249(16): p. 5176–5181.

19. Fazi, A., et al., Purification and partial characterization of the phosphoglucomutase isozymes from human placenta. Prep Biochem, 1990. 20(3-4): p. 219–40.

20. Naught, L.E. and P.A. Tipton, Formation and reorientation of glucose 1,6-bisphosphate in the PMM/PGM reaction: transient-state kinetic studies. Biochemistry, 2005. 44(18): p. 6831–6.

21. Ray, W.J., Jr. and G.A. Roscelli, A KINETIC STUDY OF THE PHOSPHOGLUCOMUTASE PATHWAY. J Biol Chem, 1964. 239: p. 1228–36.

22. Jolly, L., et al., Autophosphorylation of Phosphoglucosamine Mutase from <i>Escherichia coli</i>. Journal of Bacteriology, 2000. 182(5): p. 1280–1285.

23. Morava, E., et al., Impaired glucose-1,6-biphosphate production due to bi-allelic PGM2L1 mutations is associated with a neurodevelopmental disorder. Am J Hum Genet, 2021. 108(6): p. 1151–1160.

24. Leloir, L.F., R.E. Trucco, and et al., The formation of glucose diphosphate by Escherichia coli. Arch Biochem, 1949. 24(1): p. 65–74.

25. Veiga-da-Cunha, M., et al., Mammalian phosphomannomutase PMM1 is the brain IMP-sensitive glucose-1,6-bisphosphatase. The Journal of biological chemistry, 2008. 283(49): p. 33988–33993.

26. Maino, V.C. and F.E. Young, Regulation of glucosylation of teichoic acid. II. Partial characterization of phosphoglucomutase in Bacillus subtilis 168. J Biol Chem, 1974. 249(16): p. 5176–81.

27. Waygood, E.B., J.S. Mort, and B.D. Sanwal, The control of pyruvate kinase of Escherichia coli. Binding of substrate and allosteric effectors to the enzyme activated by fructose 1,6-bisphosphate. Biochemistry, 1976. 15(2): p. 277–282.

28. Zhang, Y., et al., The Brucella melitensis M5-90 phosphoglucomutase (PGM) mutant is attenuated and confers protection against wild-type challenge in BALB/c mice. World Journal of Microbiology and Biotechnology, 2016. 32(4): p. 58.

29. Gibson, D.G., et al., Enzymatic assembly of DNA molecules up to several hundred kilobases. Nature Methods, 2009. 6(5): p. 343–345.

30. Bertani, G., Studies on lysogenesis. I. The mode of phage liberation by lysogenic Escherichia coli. J Bacteriol, 1951. 62(3): p. 293–300.

31. Letunic, I. and P. Bork, Interactive Tree Of Life (iTOL) v5: an online tool for phylogenetic tree display and annotation. Nucleic Acids Research, 2021. 49(W1): p. W293–W296.

32. Madeira, F., et al., The EMBL-EBI search and sequence analysis tools APIs in 2019. Nucleic acids research, 2019. 47(W1): p. W636–W641.

