## Supplementary Material for "Glucose-1,6-bisphosphate, a key metabolic regulator, is synthesized by a distinct family of α-phosphohexomutases widely distributed in prokaryotes" (from another version of this preprint: version 2, 2021-12-05)

| **Name** | **Typical length (aa)** | **Sugar-binding domain sequence motif** |
|---|---|---|
| cd03085 (PGM1) | ~550 | **CGEESFG** |
| cd05799 (PGM2) | 550 – 650 | AF**EE**AIG |
| cd03089 (PMM/PGM) | ~450 | A**GEMS**G**H** |
| cd03087 (PGM_like1) | ~450 | G**GE**G**NG**G |
| cd05800 (PGM_like2) | ~450 | G**GEESGG** |
| cd05801 (PGM_like3) | ~550 | G**GEESAG** |
| cd05803 (PGM_like4) | ~450 | **GGEGNGG** |

**Table S1**: **PGM and "PGM like" subfamilies according to CDD:**

Given is the typical length and characteristic sugar-binding sequence motif of each PGM subfamily. Highly conserved amino acids are highlighted in bold

| | **cd03085** | **cd05799** | **cd03089** | **cd03087** | **cd05800** | **cd05801** | **cd05803** |
|---|---|---|---|---|---|---|---|
| *Mycobacterium tuberculosis* | | + | + | | | + | |
| *Escherichia coli* | | | + | | | + | |
| *Helicobacter pylori* | | | + | | | | |
| *Bordetella pertussis* | | | + | | | | |
| *Neisseria gonorrhoeae* | | | + | | | | |
| *Eggerthella lenta* | | | | | + | | |
| *Bacteroides salyersisae* | | + | | | + | | + |
| *Synechocystis sp. PCC6803* | | | | | + | + | |
| *Prochlorococcus marinus* | + | | | | + | | |
| *Nostoc punctiforme* | + | | | | + | | |
| *Deinococcus radiodurans* | | | | | + | + | |
| *Geobacter metallireducens* | | | | | + | | |
| *Litorilinea aerophilia* | | | | | + | + | |
| *Prosthecochloris vibrioformis* | | | | | + | + | + |
| *Granulicella mallensis* | | | | | + | + | |
| *Ammonifex degensii* | | | | + | + | | + |
| *Vibrio cholerae* | | + | | | + | + | |
| *Caldicellulosiruptor owensis* | | | | | + | | |
| *Thermotoga caldifontis* | | | | | + | | |
| *Haloplanus aerogenes* | | | | + | + | | |
| *Persephonella atlantica* | | | | | + | | |
| *Bacillus subtilis* | | + | | | | | |
| *Staphylococcus aureus* | + | + | | | | | |

**Table S2: Distribution of PHM subfamilies considered to be PGM or "PGM_like" in different bacterial strains**

Blue color with (+) shows the presence of the respective αPHM subfamily in the listed strains

| Primer | Sequence (5’ - 3’) |
|---|---|
| pASK-C(slr1334)_fw | TAGAAATAATTTTGTTTAACTTTAAGAAGGAGATATACAAATGGTTTACACTCCTGCTCC |
| pASK-C(slr1334)_rev | GGTCTTATTTTTCGAACTGCGGGTGGCTCCAGCTAGCCATGTGAGATGATTGTGCAGGCTTG |
| pASK-C(sll0726)_fw | TAGAAATAATTTTGTTTAACTTTAAGAAGGAGATATACAAATGACAAGCAGAATTAATCC |
| pASK-C(sll0726)_rev | GGTCTTATTTTTCGAACTGCGGGTGGCTCCAGCTAGCCATGCCCAAAGCCGAGGTAACAATG |
| pASK-IBA5_Cterm_fw | GGCTAGCTGGAGCCACCCGCAGTTCGAAAAATAAGACCATGGTCTCTGATATCTAACTAAG |
| pASK-IBA5_Cterm_rev | TTGTATATCTCCTTCTTAAAGTTAAAC |

**Table S3: List of the primers used in this study**

| Plasmid | Purpose |
|---|---|
| pASK-C(sll0726) | Expression of Strep-tagged Sll0726 in *E.coli* |
| pASK-C(slr1334) | Expression of Strep-tagged Slr1334 in *E.coli* |
| pET15b(PGK) | Expression of His-tagged Phosphoglycerate-kinase in *E.coli* |

**Table S4. List of the plasmids used in this study**

| Strain | Genotype | Purpose | Manufacturer |
|---|---|---|---|
| *E. coli* NEB10β | Δ(ara-leu) 7697 araD139 fhuAΔlacX74 galK16 galE15 e14-Φ80dlacZΔM15 recA1 relA1 endA1 nupGrpsL(StrR) rphspoT1 Δ(mrr-hsdRMS-mcrBC) | Molecular cloning | New England Biolabs (Ipswich (Massachusetts), USA) |
| *E. coli Rosetta gami (DE3)* | Δ(ara-leu)7697 ΔlacX74 ΔphoA PvuII phoR araD139 ahpC galE galK rpsL (DE3) F'[$lac^+$ $lacI^q$ pro] gor522::Tn10 trxB pLysSRARE ($Cam^R$, $Str^R$, $Tet^R$) | Protein overexpression | Merck Millipore (Billerica (Massachusetts), USA) |

**Table S5. List of the bacterial strains used in this study**

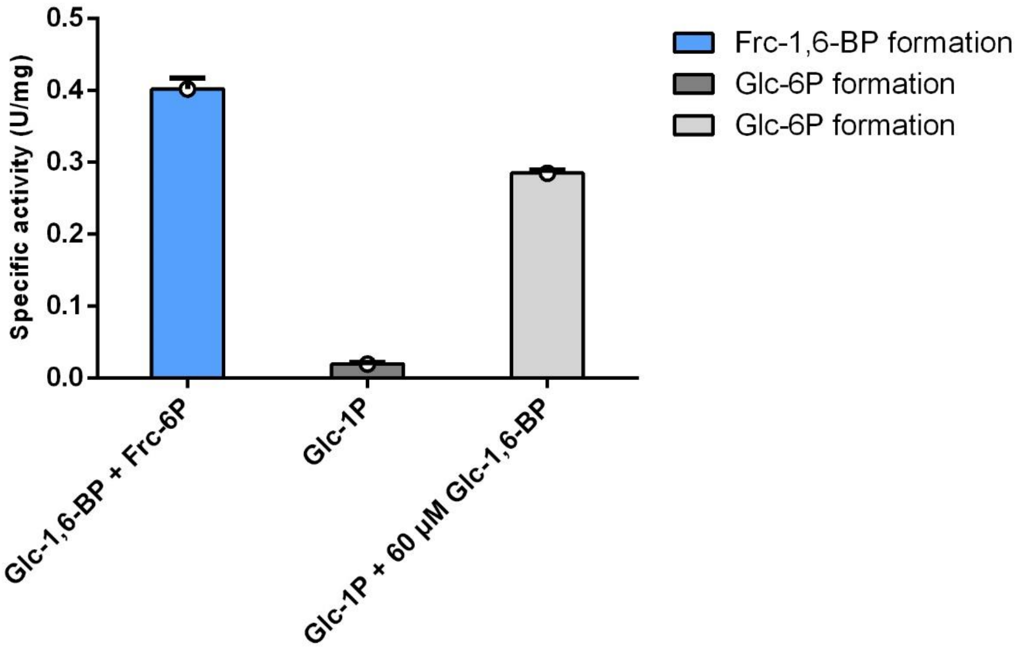


**Figure S1: Specific activity of the Slr1334 orthologue from *Bacteroides salyersiae***

Shown is: in blue, the reversible Glc-1,6-BP synthase activity of the *B. salyersiae* Slr1334 orthologue in the presence of 50 µM Glc-1,6-BP and 5 mM Frc-6P yielding Frc-1,6-BP; in dark grey and light grey, the PGM activity of the *B. salyersiae* Slr1334 orthologue in the sole presence of 1 mM Glc-1P, or in the simultaneous presence of 60 µM Glc-1,6-BP and 1 mM Glc-1P, respectively.
